# Exact Reconstruction of Gene Regulatory Networks using Compressive Sensing

**DOI:** 10.1101/004242

**Authors:** Young Hwan Chang, Joe W. Gray, Claire Tomlin

## Abstract

**Background:** We consider the problem of reconstructing a gene regulatory network structure from limited time series gene expression data, without any a priori knowledge of connectivity. We assume that the network is sparse, meaning the connectivity among genes is much less than full connectivity. We develop a method for network reconstruction based on compressive sensing, which takes advantage of the network’s sparseness.

**Results:** For the case in which all genes are accessible for measurement, and there is no measurement noise, we show that our method can be used to exactly reconstruct the network. For the more general problem, in which hidden genes exist and all measurements are contaminated by noise, we show that our method leads to reliable reconstruction. In both cases, coherence of the model is used to assess the ability to reconstruct the network and to design new experiments. For each problem, a set of numerical examples is presented.

**Conclusions:** The method provides a guarantee on how well the inferred graph structure represents the underlying system, reveals deficiencies in the data and model, and suggests experimental directions to remedy the deficiencies.

## Introduction

Mathematical modeling of biological signaling pathways can provide an intuitive understanding of their behavior [1] [2] [3]. However, since typically only incomplete knowledge of the network structure exists and the system dynamics is known to be sufficiently complex, the challenge has become to show that the identified networks and corresponding mathematical models are enough to adequately represent the underlying system. In the last years, many data-driven mathematical tools have been developed and applied to reconstruct graph representations of gene regulatory networks (GRNs) from data. These include Bayesian networks, regression, correlation, mutual information and system-based approaches [4] [5] [6] [7] [8] [9] [10]. Also, these approaches either focus on static or on time series data. The latter approach has the advantage of being able to identify dynamic relationships between genes.

However, data-driven reconstruction of the network structure itself remains in general a difficult problem; nonlinearities in the system dynamics and measurement noise make this problem even more challenging. For linear time invariant (LTI) systems, there exist necessary and sufficient conditions for network reconstruction [11]. However, for time-varying or nonlinear systems, there has not been as yet any statistical guarantee on how well the inferred model represents the underlying system [12] [13] [14] [15]. The recent work [16] addresses the problem of data-driven network reconstruction, together with measurement noise and unmodelled nonlinear dynamics, yet this work points out that these complications impose a limit on the reconstruction, and with strong nonlinear terms the method fails. Additionally, identifying whether important nodes in the graph structure are missing, how many are missing, and where these nodes are in the interconnection structure remains a challenging problem.

Moreover, in order to continue to have an impact in systems biology, identification of the graph topology from data should be able to reveal deficiencies in the model and suggest new experimental directions [17]. For example, Steinke *et al.* [18] presented a Bayesian method for identifying a gene regulatory network from micro-array measurements in perturbation experiments and showed how to use optimal design to minimize the number of measurements.

Since biological regulatory networks are known to be sparse, meaning that most proteins interact with only a small number of proteins compared with the total number in the network, many methods [12] [13] [14] [15] [19] [20] [21] [22] take advantage of the sparsity. The methods typically use *l*_1_-norm optimization, which leads to a sparse representation of the network and improves the ability to find the actual network structure. Even though many methods [14] [15] show that the reconstruction results are fairly good, the methods cannot guarantee *exact recovery*. This stems from the fact that the so-called *incoherence* condition is typically not satisfied for the matrix **Ω** in a linear measurement model **Y = Ωq**, where **q** is the signal to be reconstructed, **Y** is the measurement, and **Ω** is known as the sensing matrix. Roughly, incoherence is a measure of the correlation between columns of the sensing matrix. Since the incoherence condition of the sensing matrix provides a metric of performance, this is one of the motivating factors for the use of compressive sensing (CS) [23]. CS is a signal processing technique for efficiently acquiring and reconstructing a signal by taking advantage of the signal's sparsity and allowing the entire signal to be determined from relatively few measurements under a certain condition, i.e., it requires that the incoherence condition to be satisfied. In the Human Epidermal Growth Factor Receptor2 (HER2) positive breast cancer signaling pathway that we studied in [24] [25], time series data sets consist of only 8 time point measurements of 20 protein signals, and we would like to use this limited data to identify a graph structure which could have 20 x 20 or 400 edges.

In this paper, we are interested in directed graph representations of signaling pathways. We develop a new algorithm for GRN reconstruction based on CS. First, we focus on sparse graph structures using limited time series data with all nodes accessible and no measurement noise. We test the network reconstruction algorithm on a simulated biological pathway in which the structure is known *a priori.* We explain the limitation of the proposed method’s performance when the dataset naturally has high coherence and propose a way to overcome this limitation by designing additional effective experiments.

Next, the proposed algorithm is extended to a more general problem: we consider partially corrupted data with data inconsistencies due to model mismatch, and measurement noise affecting all the data. Typically, data inconsistencies may result from missing nodes in the model; or in some cases arbitrary data corruption may result from human errors such as mislabeling or the improper use of markers or antibodies. The question is whether one can still recover the graph structure reliably under these conditions. Inspired by a robust error correction method [26], the exact recovery of the graph structure can be guaranteed under suitable conditions on the node dynamics, provided that hidden nodes can affect relatively few nodes in the graph structure. Also, a set of numerical examples is provided to demonstrate the method, including some from an RPPA (Reverse Phase Protein Array) dataset [27] collected from HER2 positive breast cancer cell lines. In this paper, the main contributions are the following.

- The CS framework uses the coherence of the sensing matrix as a performance index, which allows us to assess and optimize mathematically network reconstruction.
- Coherence also provides a guideline for optimizing experiment design for network reconstruction.
- By utilizing an error correction method in conjunction with the CS framework, network reconstruction may be performed even when there are hidden nodes and measurement noise.

## Background

### Overview: Compressive Sensing

Consider measurements Y ∈ ℝ^*m*^ of a signal q ∈ ℝ^*n*^:

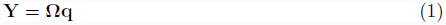

where **Ω** ∈ ℝ^*m*×*n*^ is called the sensing matrix.

One key question [28] is how many measurements *m* are needed to exactly recover the original signal q from **Ω**:

- If *m* > *n* and **Ω** is a full rank matrix, then the problem is overdetermined. If *m* = *n* and **Ω** is a full rank matrix, the problem is determined and may be solved uniquely for q.
- If *m* < *n*, the problem is underdetermined even if **Ω** has full rank. We can restrict q ∈ ℝ^*n*^ to the subspace which satisfies **Y = **Ω**q**. However, q cannot be determined uniquely.

For the underdetermined case, the least squares solution 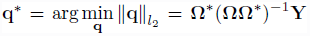 is typically used as the “best guess” in many applications. However, if q is known to be sparse, meaning that many of its components are zero, one might expect that fewer than *n* measurements are needed to recover it. It is thus of interest to obtain a good estimator for underdetermined problems in which q is assumed to be *s-sparse*, meaning that s of the elements of q are nonzero. In principle, the theory of compressive sensing (CS) asserts that the number of measurements needed to recover q ∈ ℝ^*n*^ is a number proportional to the compressed size of the signal (s), rather than the uncompressed size (*n*) [29]. To be able to recover **q**, CS relies on two properties: *sparsity*, which pertains to the signals of interest, and *incoherence*, which pertains to the sensing matrix.

#### Proposition 1

*[28]*.. *Suppose that any 2s columns of an m* ×*n matrix* **Ω** *are linearly independent (this is a reasonable assumption if m*≥ *2s*). *Then, any s-sparse signal* **q** ∈ ℝ^*n*^ *can be reconstructed uniquely from* **Ω**q.

The proof [28] of the above proposition also shows how to reconstruct an *s-sparse* signal q ∈ ℝ^*n*^ from the measurement **Y = **Ω**q** where q is the unique sparsest solution to **Y = **Ω**q**:

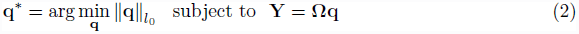

and 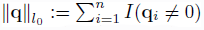 is the cardinality of q. However, the *l*_0_-minimization is computationally intractable (NP-hard in general). Recent breakthroughs enable approximating *l*_0_-optimization by using *l*_1_-minimization which is a convex optimization problem and can be solved in a simple but effective way by linear programming:

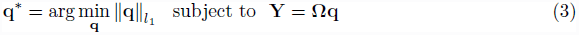

The *l*_1_-minimization (3) requires mild oversampling, more specifically, *m* ≥ *c*. *μ* (Φ, Ψ)^2^ • *s* log*n* for some positive constant c where we have m measurement in the Φ domain under bases Ψ and *μ* represents the coherence defined as follows:

#### Definition 1

*[23]*.. *The coherence of a matrix* **Ω** *is given by*

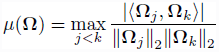

*where* **Ω**_*j*_ *and denote columns of* **Ω**.

Several theoretical results [30] ensure that the *l*_1_-minimization guarantees exact recovery whenever the sensing matrix **Ω** is sufficiently *incoherent.* For example, we can say that **Ω** is *incoherent* if μ is small. Coherence is a key property in the compressed sensing framework because if two columns are closely correlated, it would be very difficult to distinguish whether the influence from the components in the sparse signal comes from one or the other (recall that the measurement Y(= **Ω**q) is a linear combination of each columns of the sensing matrix **Ω** with the components in q as coefficients). Also, numerical experiments suggest that in practice, most s-sparse signals are in fact recovered exactly once m ≥ 4*s* [23]. Here, “exact recovery” means that we find the sparsest solution (q) such that **Y = **Ω**q**.

Therefore, if the sensing matrix **Ω** satisfies the incoherence condition (m ≥ *c*.*μ*(Φ, Ψ)^2^ •*s* log *n*, or in practice, m ≥ 4*s*), a sufficiently sparse signal q can be exactly recovered from the limited dataset without any prior knowledge of the number of nonzero elements, their locations, and their values. On the other hand, if the condition is not satisfied, exact recovery cannot be guaranteed [29] [17]. However, it is possible to use the property of coherence to guide biological experiment design, basically to collect a more informative dataset. As we will discuss in this paper, this can be done by inhibiting or stimulating certain genes to manipulate the gene expression.

## CS can help reconstruct GRNs

In graph theory, a digraph can be represented by G = (*V, E*) where *V* and *E* represent nodes and edges respectively. For GRNs, each node represents a gene and each edge represents an influence map which models how genes affect each other. For example, the interactions could be how genes inhibit or stimulate each other. Since the connectivities of GRNs are typically unknown, often the best we can do is to select a set of possible candidate functions encoding possible unknown connectivities between genes.

In this section, we formulate a data-driven network identification problem into CS framework: first, we define a dynamical model of gene regulatory network. Then, we encode system dynamics into the sensing matrix (**Ω**) and denote unknown connectivities between genes by **q**, a signal to be recovered.

### A Dynamical Model of Gene Regulatory Networks (GRNs)

We consider a dynamical system described by:

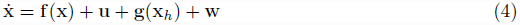

where x ∈ ℝ^*n*^ denotes the concentrations of the rate-limiting species which can be measured in experiments; 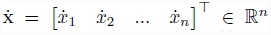 is a vector whose elements are the change in concentrations of the *n* species over time; f(.): ℝ^*n*^ → ℝ^*n*^ represents biochemical reactions, such as those governed by mass action kinetics, Michaelis-Menten, or Hill kinetics. Thus, f(.) can include functions of known form such as product of monomials, monotonically increasing or decreasing Hill functions, simple linear terms, and constant terms [19]. u ∈ ℝ^*n*^ denotes the control input which could represent inhibitions and stimulations; g(.): ℝ^*nh*^ → ℝ^*n*^ represents influence from hidden nodes x_*h*_ ∈ ℝ^*nh*^, which cannot be measured in experiments; n_*h*_ is the number of hidden nodes and unknown; and w ∈ ℝ^*n*^ represents energy-bounded process noise or measurement noise. Here x, X which can be can be measured in experiments, and u is assumed to be known.

Since we do not know whether important nodes in the gene regulatory network are missing, how many missing nodes there are, and how they affect system dynamics (i.e., x_*h*_, *n*_*h*_ and g(x_*h*_)), we denote a vector of (unknown) influence from hidden nodes’ dynamics by v(≜ g(x_*h*_)); Also, without loss of generality, since we know u, we define y as follows:

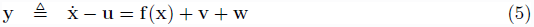

### Formulating a Dynamical System as a GRN

The nonlinear function f(x) can be decomposed into a linear sum of scalar basis functions, *f*_*b,i*_(x) ∈ ℝ, where we select the set of possible candidate basis functions:

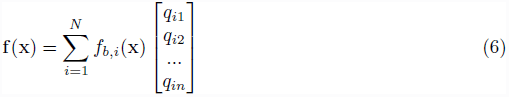

where *N* is the number of possible candidate basis functions and *q*_*ij*_ are unknown parameters which reflect underlying structure, i.e., influence of *f*_*b,i*_(**x**) on the *j*-th state 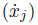. Typically, we may choose a larger set than necessary, and allow the CS method to indicate the importance of each function, as we shall describe. Thus the equation (5) can be written as follows:

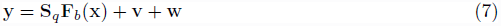

where 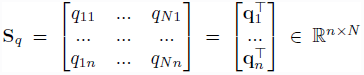 reflects the underlying GRN structure,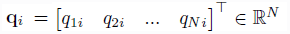 and 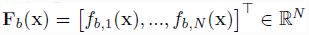 is the vector field which includes possible candidate basis functions. In this way, any biochemical reactions can be represented by a linear map S_*q*_ and a function F_***b***_(x) where S_*q*_ encodes the underlying graph structure and F_***b***_(x) includes all possible candidate functions that could be included in the biochemical reactions.

In practice, we can construct F_***b***_(x) by selecting the most commonly used candidate basis functions to model GRNs, for example, all monomials, binomials, other combinations and Hill function.

#### Example 1.

*Consider the simple nonlinear ordinary differential equations (ODEs):*

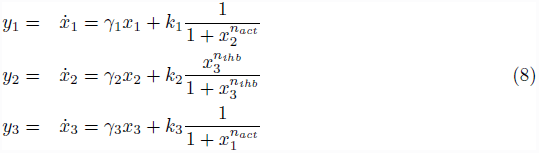

**Figure 1.**
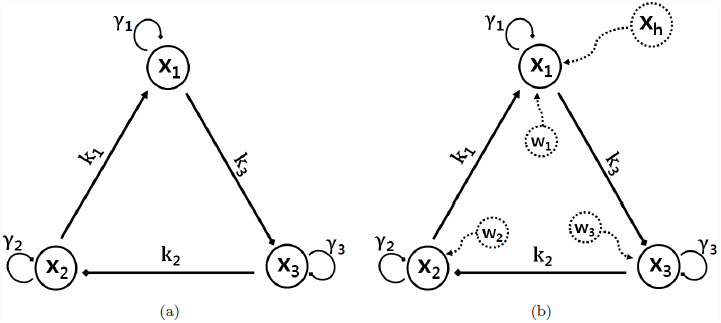
A graph representation of nonlinear ODEs. (a) (Example 1, *n* = 3, S_*q*_ ∈ ℝ^3x6^): among 18(= 3 × 6) components, only 6 components are non-zero (b) (Example 1 with hidden node and measurement noise) there exists hidden node x_*h*_ affecting x_1_ and process noise **w**.

*where x_*i*_ denotes the concentration of the i-th species, 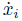 is the change in concentration of the i-th species, the γ_*i*_ denotes protein decay rate, the k_*i*_ denotes the maximum promoter/inhibitor strength. Here, there is no input* (**u = 0**), *no hidden node* (**v = 0**) *and no process noise* (**w = 0**). *Also, n_act_ represents positively cooperative binding (activation) and n_ihb_ represents negative cooperative binding (inhibition). The set of ODEs corresponds to a topology where gene 1 is activated by gene 2, gene 2 is inhibited by gene 3, and gene 3 is activated by gene 1 as shown in Figure 1 (a). We can write (8) as follows:*

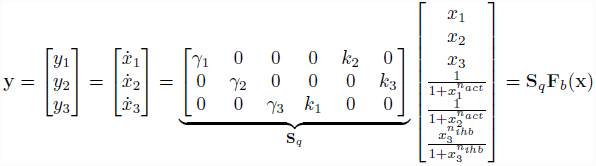

*where* S_*q*_ ∈ ℝ ^3×6^ *represents the influence map.*

*We can also consider a version of (8) in which there exists a hidden node (x_h_) affecting (x*_1_*) as shown in Figure 1(b), as well as process noise.*

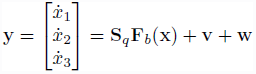

*where* v = [*v*_1_, 0, 0]^T^ *and* w = [*w*_1_, *w*_2_, *w*_3_] ^T^

### Formulating GRN into the CS framework

Suppose the time series data are sampled from a real experimental system at discrete time points *t*_*k*_. By taking the transpose of both sides of equation (7), we obtain

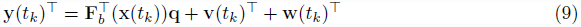

where 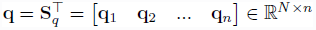.Assuming thatM successive data points are sampled,then define:

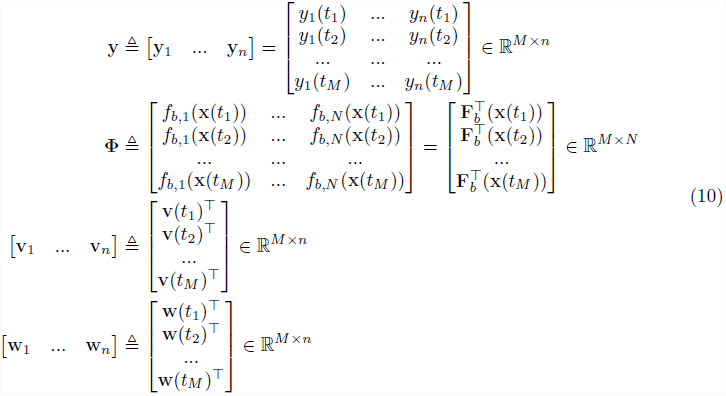

This leads to *n* independent equations:

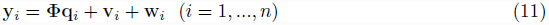

where 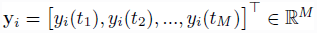 represents the *M* successive data points, Φ ∈ ℝ^*M*×*N*^ consists of *N* possible candidate bases which are functions of given time series data x and q_*i*_ ∈ ℝ^*N*^ represents the unknown influence map corresponding to the *i*-th species. Since a biochemical reaction network is typically sparse, as a consequence, q*_i_* is sparse and we have *N* ≫ *M* for Φ because we assume the limited time series data and may choose a larger set of basis functions than necessary.

Although we formulate *n* independent linear regression problems (11), we consider *n* independent equations in (11) together by stacking y*_i_* as follows:

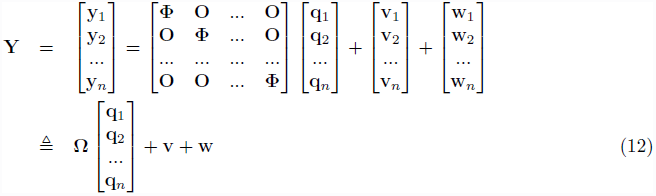

Now, equation (12) is in the form of the CS formulation in (1) where Y ∈ ℝ^*n.M*^ is the measurement, **Ω** ∈ ℝ^*n.M*×*n.N*^ is the sensing matrix which consists of basis functions for the given time series data,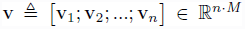 represents possibly large corruption from hidden nodes and 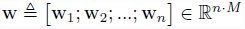 represents process or measurement noise.

In this paper, we make the following assumptions:

#### Assumption 1.

*We consider two cases:*

- *(ideal case) We assume that we can measure all states* x *(i.e., there is no hidden node* (x_*h*_)) *and there is no measurement noise* (w = 0). *Also, there are enough columns in* Φ *in (11) to represent the underlying system* (6).
- *(extension) We consider hidden node* (x_h_) *where* v(= g(x_*h*_)) *is assumed to be sparse, and process noise* (w ≠0). *Here, we can also consider the case where the columns (also known as the dictionaries in the CS literature [23]) of* Φ *may not be able to represent the underlying system. In this case, we consider the influence from the missing dictionaries as* v.

We first consider the ideal case for simplicity in explaining the main results, and then extend the proposed method to the more general case.

## Results and Discussion

### 1. Formulating GRN Identification Problem into CS

Many existing algorithms [16] [19] consider the *n* independent linear regression problems (11) separately. Since the columns of matrix Φ are composed of time series data as in (10), it is difficult to *a priori* guarantee low correlation and sometimes Φ even suffers rank deficiency. Also, Pan *et al.* [19] pointed out that correlation between the columns of Φ is usually high (μ(Φ) is close to 1).

Intuitively, if two columns of the sensing matrix are highly correlated, it is hard to distinguish the corresponding components in the sparse signal q (note that the measurement Y is a linear combination of each column of the sensing matrix with the components in q as coefficients). In order to deal with high coherence in (11), many methods combine CS with different techniques such as Bayesian formulation [19], Kalman filter [20], and Granger causality [21]. Also, it is a well known problem in the lasso formulation of network inference: if there is high coherence in the sensing matrix, one can use an elastic net which combines the *l*_1_ and *l*_2_ norms. Although each reconstruction result [19] [20] [21] might be the optimal solution in the sense of its formulation (i.e., maximum likelihood), the identified graph may not represent the underlying GRN. In other words, if the data set is not informative enough to fully explore the underlying system, while the identified graph structure based on the given data set may be an optimal solution of the particular optimization problem, it many not represent the true system.

In this paper, our goal is to get the smallest data informative enough to recover the underlying graph structure exactly. Since the proposed method maintains the CS framework by reducing coherence of the sensing matrix, the method is fundamentally different than any other methods which make use of different techniques in conjunction with the *l*_1_ optimization [19] [20] [21] which leads to a sparse representation of the network. Hence, we use all the properties of CS in order to access the ability to exactly reconstruct the underlying graph structure, reveal deficiencies in the data and model, and design new experiments to remedy the deficiencies if necessary.

While maintaining the CS framework, in order to deal with high coherence, we formulate (12) instead of (11) and we have strongly uncorrelated columns in **Ω**. In other words, since **Ω** has many independent columns, we have more degrees of freedom to reduce coherence of **Ω**. We will show that by using a transformation, the components of the sensing matrix can be made more uniformly distributed so that we could reduce coherence.

Moreover, since each q_*i*_ has different degrees of sparsity in general, if we consider *n* independent equations in (11), Φ should satisfy the incoherence condition *M* ≥ 2max_*i*_(||q_*i*_||_0_) stated in Proposition 1. On the other hand, **Ω** only needs to satisfy the condition 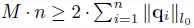. Since the averaged sparsity 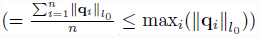 is smaller than max_*i*_(||q_*i*_||_*l*0_), we can reduce the required number of samples (*M*). Also, in case of rank deficiency, we can simply remove the corresponding rows (note that the rank deficiency is more likely caused by the row since *N* » *M*).

### 2. Optimal Design of Sensing Matrix

#### 2.1. Reducing Coherence by Transformation

In order to reduce coherence, first we rearrange **Ω** with respect to a spatial information, and then we consider the transformation in order to reduce coherence (see Method: 1.1. Rearranging the sensing matrix for mathematical details):

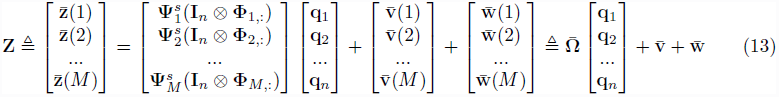

where ⊗ represents the Kronecker product ^1^, **Ω** ∈ ℝ^*M*.*n×N*.*n*^ represents the rearranged sensing matrix multiplied by the transformation 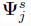, and 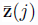, 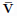 and 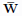 are defined in (20). We want to find 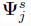(*j* = 1, … *M*) to minimize where

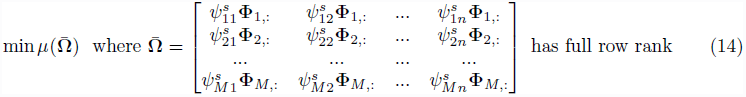

where 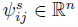represents the *j*-th column of 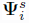. In this paper, we propose a heuristic approach and a novel way to find 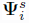 by solving the optimization iteratively to reduce coherence (see Method: Section 1.2 and 1.3 for details).

**Figure 2.**
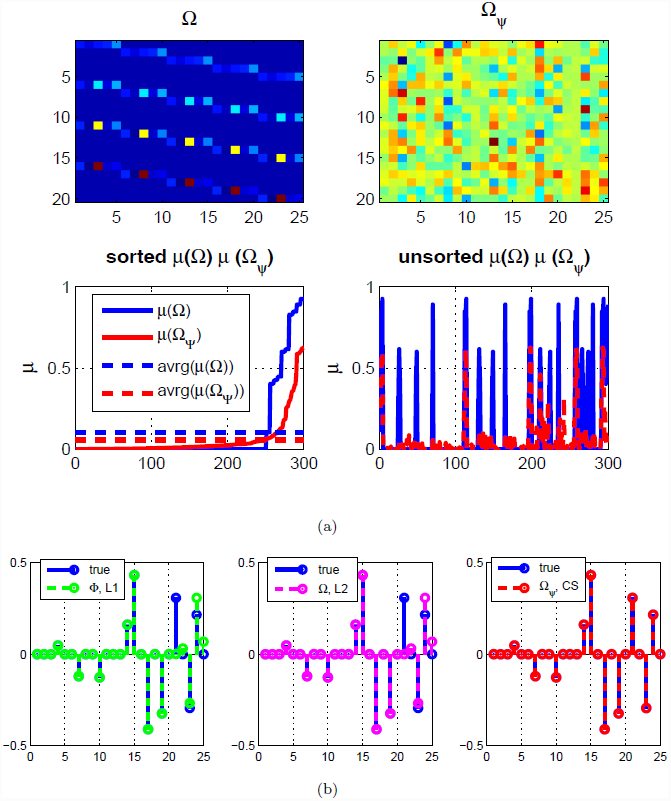
Reducing coherence for a linear system. (Example 2) (a) the sensing matrix **Ω**, without and with transformation Ψ (top) and coherence (bottom) (b) reconstruction result (left) *L*1 (middle) *L*2 with **Ω** (right) *l*_2_ optimization with **Ω**_Ψ_(CS).

##### Example 2.

*(reducing coherence for a linear system) consider a simple linear system* 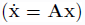 *where n* = 5, *N* = 5, *M* = 4, *s* = 10 *(note that n. M ≥ 2s), and the elements of* **A** *are randomly chosen such that there is no isolated node. Figure 2 shows the sensing matrix for both* **Ω** *(top left) and xthe transformed sensing matrix (top right, denoted by* **Ω**_ψ_). *By reducing coherence, the components of the transformed sensing matrix (top right) are more uniformly distributed and the coherence is reduced by up to* 0.6 *although* μ(**Ω**) *is close to 1 (bottom left) in Figure 2(a). Also, Figure 2(b) shows the result of the inferred graph structure based on given time series data without any a priori information where the x-axis represents indices of the influence map (i.e., the* 1_*st*_, 2_*nd*_, …,n_*th*_ rows of influence map S_*q*_; note that for a linear system, **S_*q*_ = A**). *Here, there are 5 states (n* = 5) *in a linear system so the influence map* A *has 25 elements. Although L1 and L*2^2^ *norm minimizations fail to recover the exact signal, CS in (17)* (**see 3. Recovery of Gene Regulatory Networks** *for details) recovers the exact signal (bottom right) by first reducing coherence of the sensing matrix. Note that* (L1) *solves the n independent equations (11) without reducing coherence and* (L2) *solves (12) with l_2_-regularization*.

##### Example 3.

*(reducing coherence for a nonlinear system*, *n* = 3, *N* = 9, *M* = 5, *s* = 6) consider simple nonlinear ODEs as follows:

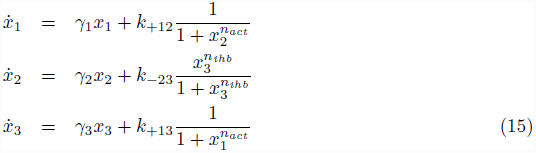

**Figure 3.**
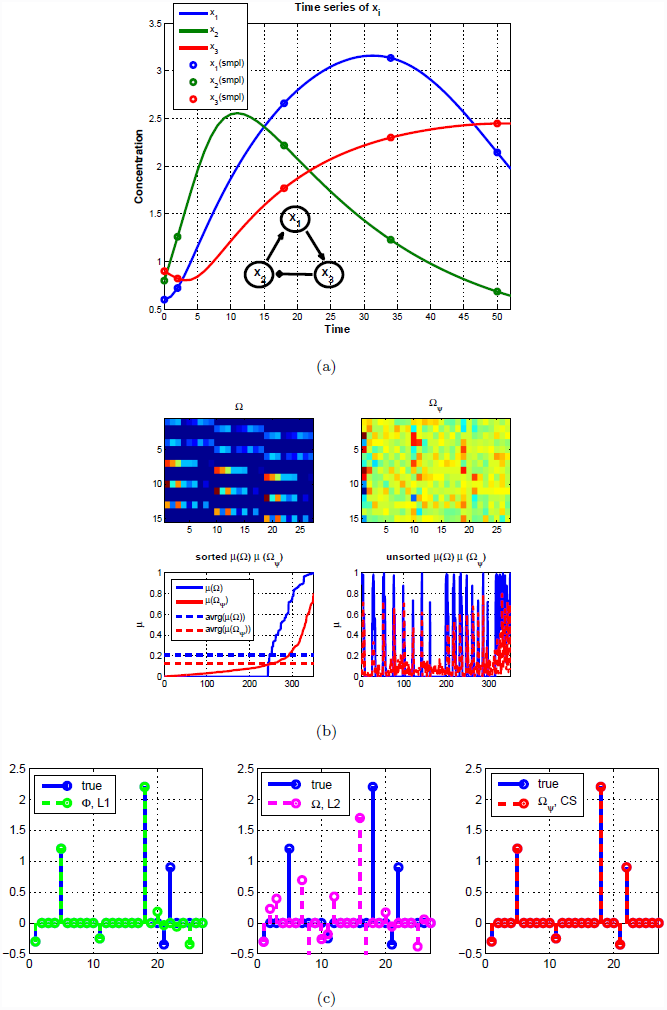
Reducing coherence for a nonlinear system. (Example 3) (a) Time series of *x*_1_, *x*_2_, *x_3_* for model (15) (b) the sensing matrix w/o, and w/ Ψ matrix (top) and coherence (bottom) (c) reconstruction results.

*where*(*γ*_1_,*γ*_2_,*γ*_3_) = (−0.3, −0.25, −0.35), k_+12_ = 1.2, k_+13_ = 0.9, *k_-23_* = 2.2, *n_act_* = 4 *(activation Hill coefficient) and n_ihb_* = 4 *(inhibition Hill coefficient). The set of ODEs corresponds to a topology (x_2_ → x_1_ → x_3_* ⊣ x_2_) *as shown in Figure 3 (a). Figure 3(b) shows that we can reduce the coherence by up to* 0.8 *and (c) shows that only CS recovers the exact graph structure, and *L*2 regularization does not encourage sparsity but distributes the coefficients to be more similar to each other*.

#### 2.2. Designing Effective Experiments

Consider the case where the sensing matrix is not incoherent. If the coherence condition 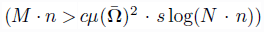 is not satisfied, exact recovery cannot be guaranteed [29]. We use the transformation (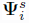; or P=*P*^T^*P* in (24); see Method section 1. Reducing Coherence by Transformation) in order to reduce the coherence but obviously, sometimes we might have inherent limits to how much the coherence can be reduced. There are possible reasons:

- Since we solve the relaxed problem (24) iteratively, P might be sub-optimal.
- If the time series data of two different gene expressions, x_*i*_ and x_*j*_ are highly correlated, it might be difficult to reduce coherence. In this case, we need to design a new experiment to remedy deficiencies in the data.

As we mentioned, the incoherence of the sensing matrix can be used not only as a good metric to guarantee exact recovery but also as a guideline for designing new experiments. For example, from the coherence distribution, we can identify which columns of the sensing matrix have high coherence, i.e., *f*_*b*,*i*_(x) and *f*_*b*,*j*_(x) in (6). Intuitively, in order to reduce ambiguities from the highly correlated columns of the sensing matrix, we should perturb either *f*_*b,i*_(x) or *f*_*b,j*_(x). Thus, it is possible to use this property of coherence to guide biological experiment design, to collect a more informative dataset.

**Figure 4.**
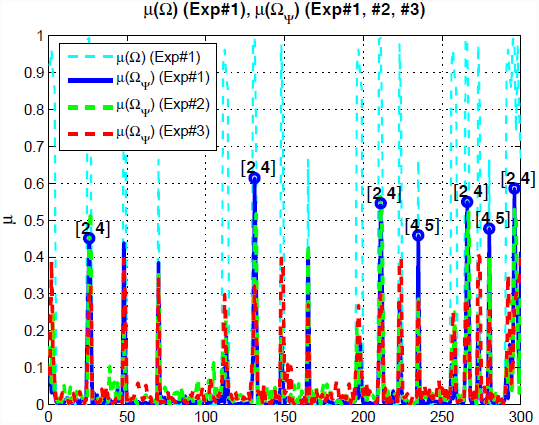
Guideline for designing new experiment based on coherence distribution. (Example 4) coherence distributions of 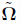 show that *x*_2_ and *x*_4_ cause high coherence.

##### Example 4.

*(limitation of reducing coherence by* Ψ, *linear dynamics, n* = 5, *N* = 5, *M* = 4, *s* = 10*) In Figure 4, Exp#1 represents the original experimental data set which has the limitation of reducing coherence by* Ψ. *Since we consider linear dynamics, i.e., f_b,i_(x)* = *x_i_ and f_b,j_*(*X*) = *x*_*j*_, *we found that x*_2_ *and x*_4_ *cause high coherence as shown in Figure 4 (circle marker). Thus, in order to reduce this high coherence, we should perturb either x*_2_ *or x*_4_.

*To show the effectiveness of the new experiment, we design two different experiment sets and compare the reconstruction results with each other; for Exp#2, we perturb x*_3_ *and for Exp#3, we perturb x*_2_. *As we mentioned earlier, intuitively, we expect that Exp#3 to be a more informative experiment to identify the graph structure since we would like to reduce the coherence between x*_2_ *and x*_4_. *As we expected, in Exp#3, we can reduce the coherence more than that of Exp#2 as shown in Figure 4, and recover the exact graph structure as shown in Figure 5. On the other hand, the coherence of Exp #2 remains almost the same as that of Exp #1 and we fail to recover the exact graph structure. This numerical example illustrates that by using the property of coherence, we can guide biological experiment design more effectively*.

**Figure 5.**
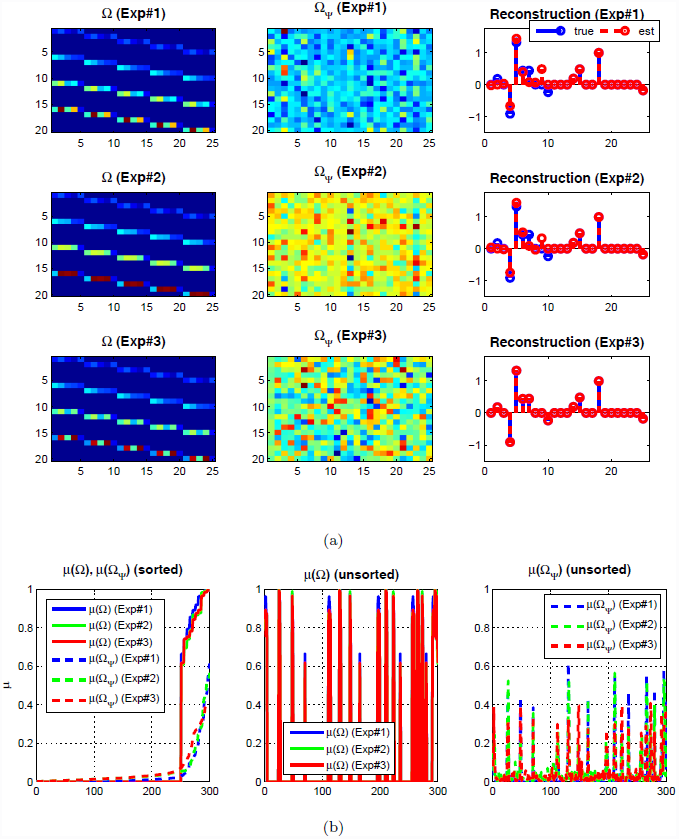
Design effective experiment for a linear system. (Example 4) Reconstruction results based on different experiments: Exp#1 represents the original experimental dataset which has limitation to reduce coherence; Exp#2 represents non-effective experimental dataset; and Exp#3 represents the effective experimental dataset (a) the sensing matrix and reconstruction result (b) coherence comparison.

##### Example 5.

(*limitation of reducing coherence by Ψ, nonlinear dynamics*, *n* = 3, *N* = 9, *M* = 5, *s* = 9) *consider the following set of ODEs*:

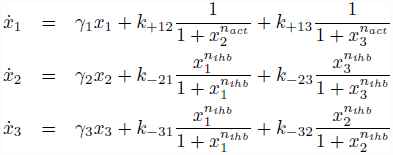

**Figure 6.**
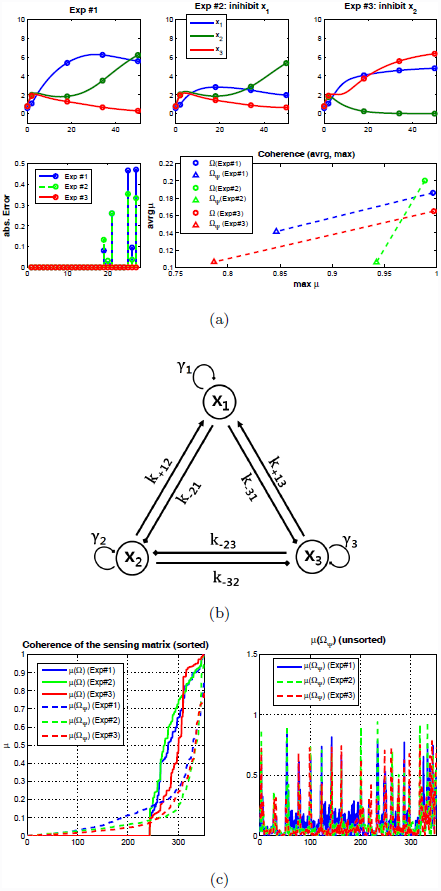
Design effective experiment for a nonlinear system. (Example 5) Reconstruction result based on different experiment (nonlinear case): Exp#1 represents the original experimental dataset which has limitation to reduce coherence; Exp#2 represents non-effective experimental dataset (inhibit *x*_1_); and Exp#3 represents the effective experimental dataset (inhibit *x*_2_) (a) the time series of *x*_1_, *x*_2_, *x*_3_ for each experiment and coherence comparison for each experiment (b) the corresponding topology (c) (detail) coherence of the sensing matrix.

*where* (γ_1_,γ_2_,γ_3_) = (−0.25, −0.23, −0.26), *k*_+12_ = 1.2, *k*_+13_ = 1.25, *k*_−21_ = 2.8, *k*_−23_ = 2.1, *k*_−31_ = 2.7, *k*_−32_ = 1.8, *n*_*act*_ = 4 *(activation Hill coefficient) and n _ihb_* = 4 *(inhibition Hill coefficient). The corresponding topology is shown in Figure 6 (b) (note that we intentionally choose the symmetric structure and similar parameters). The reconstruction error using Exp#1 data is shown in Figure 6 (a) (left, bottom) and the reconstruction error illustrates difficulties of resolving ambiguities from x*_2_ *and x*_3_. *This can be captured by the coherence distribution of the sensing matrix based on the Exp#1 dataset; the correlation between the columns corresponding to x*_1_ *and to x*_2_ *is close to the correlation between the columns corresponding to x*_1_ *and to* x_3_. *Based on the coherence distribution, we design two trials; for Exp#2, we perturb x*_1_ *and for Exp#3, we perturb x*_2_. *As we expected, Exp #2 is not an effective experiment in terms of information. On the other hand, by using Exp#3, we can reduce both the maximum coherence and the averaged coherence, and reconstruct the exact graph structure as shown in Figure 6 (a)*.

Both Examples 4 and 5 illustrate that if the transformed sensing matrix is not incoherent enough to guarantee exact recovery, we can design a new experiment based on the distribution of coherence. Also, we show that an informative experiment can help to reduce coherence and thus reconstruct the exact graph structure. For a fair comparison, we use the same number of time points as M here. However, in practice, we can stack all the experimental data sets together if we assume that the linear map S_*q*_ and the set of basis functions F_*b*_(x) does not change for different experiment:

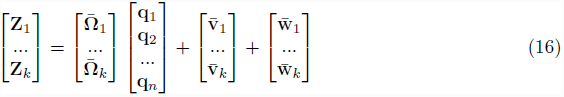

where the subscript Z*_i_* 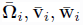 represents the *i-*th experiment. As the number of measurements increase (*M*), one may be able to reduce the coherence. However, one can reduce the coherence only if the additional measurements provide us more useful information. As a trivial example, one could stack exactly the same data on top of the first, and increase *M* to 2*M*,however the coherence is exactly the same as that of the original dataset.

### 3. Recovery of Gene Regulatory Networks

In this section, we present reconstruction of the exact graph structure and show how the condition for exact recovery will be used. First, we consider the ideal case where there are no hidden nodes and no measurement noise. Second, we extend the ideal case to the more general case.

#### 3.1. Reconstructing Gene Regulatory Networks (ideal case)

In (13), q represents the *s*-sparse network structure which we want to reconstruct from the time series gene expression by solving the *l*_1_-norm optimization

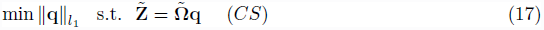

where 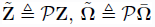 and *P* is the optimal transformation in (24)

##### Proposition 2..

*If the sensing matrix* 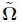 *constructed from time series data, multiplied by the optimal transformation, *P*, has 2*s* linearly independent columns, then any s-sparse network structureq can be reconstructed uniquely from 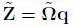*.

*Proof.* (Suppose not), then there are two *s-sparse* graph structures q_1_, q_2_ with 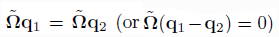.However,q_1_-q_2_ is 2*s-sparse*, so there is a linear dependence between 2s columns of 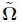 (contradiction).

The requirement of 2s linearly independent columns in **Proposition 2** may be translated to an incoherence condition on the sensing matrix. That is, if the unknown *s-sparse* signal q is reasonably sparse, it is possible to recover q under the incoherence condition on the sensing matrix. Although the sensing matrix consists of redundant dictionaries, the coherence of the sensing matrix can be reduced. In a heuristic way, we multiply the redundant dictionaries Φ_k,:_ by a randomly chosen matrix 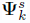 at each time step *k* and iterate this step until the coherence is decreased. Or, we can find the optimal transformation *P*(or P) in (24) to reduce the coherence. In the previous numerical examples, we illustrated that the coherence of the sensing matrix is decreased by transformation and showed the exact reconstruction of graph structure.

**Figure 7.**
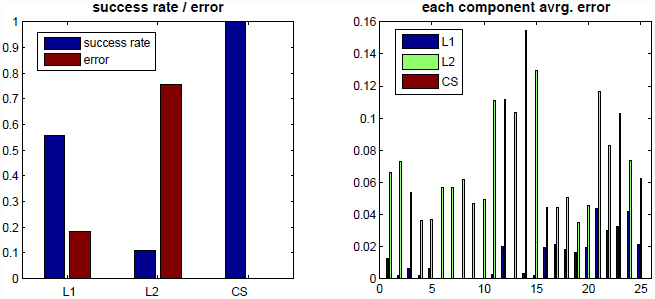
Comparison *L*1,*L*2, and *CS*. (Example 6, linear case) statistics of 50 trials: (left) success rate and reconstruction error (right) reconstruction error for each component.

##### Example 6.

*(Statistics) Here, we compare the success rate of the proposed method with other methods such as L1 and L2. Figure 7 shows statistics of* 50 *trials for a simple linear case (for each trial, we randomly generate the influence map). Here, we count the number of successes of each method when any of the methods recover the exact structure. By reducing coherence, we can improve the success rate as shown in Figure 7 (a). Also, L2-regularization does not encourage sparsity but distributes the coefficients to be more similar to each other as shown in Figure 7 (b)*.

#### 3.2. Graph Reconstruction with Hidden Nodes

The main contributions of the proposed method in the previous section is the conversion of the problem of inferring graph structure into the CS framework. Then, we demonstrate that one could recover sparse graph structures from only a few measurements. However, for practical use, the proposed method needs to be able to deal with both sparsely corrupted signals (v = g(x_*h*_)) and measurement noise (w) in (5).

In general, the assumption of accessibility or observability of all nodes [17] is not satisfied. Thus, we focus on the case in which the hidden node affects observable nodes directly as shown in Figure 1 (b). Also, without loss of generality, the hidden node dynamics could be any arbitrary dynamic model. Or, even if there is no hidden node, a small portion of the biological experiment dataset could be in practice contaminated by large error resulting from, for example, mislabeling, or improper use of markers or antibodies. Moreover, all biological datasets are contaminated by at least a small amount of noise from measurement devices. Therefore, the proposed method should be robust. We note this goes beyond the results in [11] due to the consideration of hidden node dynamics with measurement noise.

Here, the question is whether it is still possible to reconstruct the graph structure reliably when measurements are corrupted. Since hidden nodes and measurement noise are considered, the number of time points is assumed to be greater than that of the previous case [17] (i.e., no corruption and no measurement noise). Thus, the number of rows of the sensing matrix is assumed to be greater than the number of columns. If the number of the time points *M*(< *N*) is limited, then we can stack 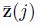 with different 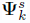 or including different dataset as described in equation (16).

In CS literature [26], two decoding strategies for recovering the signal from a corrupted measurement are introduced, where the corruption includes both a possible sparse vector of large errors and a vector of small error affecting all the entries. It is shown that two decoding schemes allow the recovery of the signal with nearly the same accuracy as if no sparse large errors occurred. Our contribution is converting the problem of inferring the graph structure with hidden nodes into the highly robust error correction method framework [26] (see Method: Section 2. Two step refinements) and showing how this can improve the reliability of reconstruction.

##### Example 7.

*(arbitrary corruption with no measurement noise) Consider nonlinear ODEs as follows:*

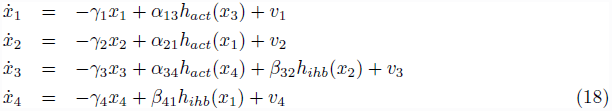

**Figure 8.**
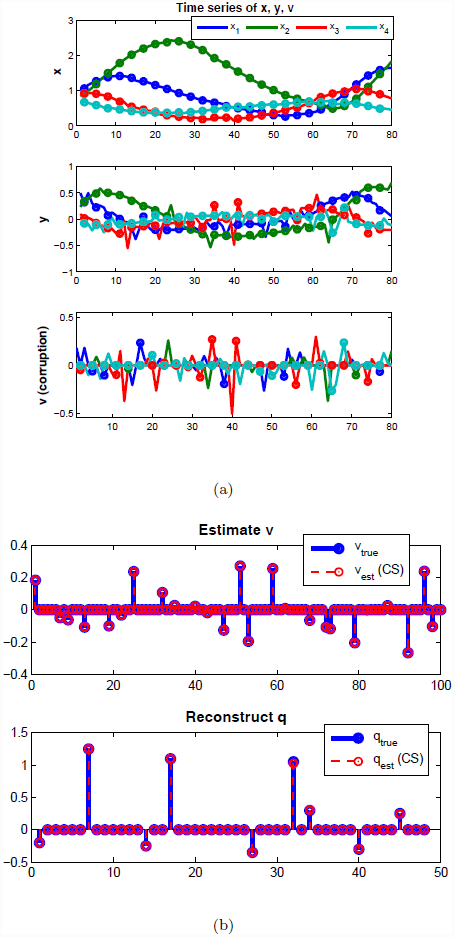
Arbitrary corruption with no measurement noise. (Example 7) Reconstruction with corrupted signal. (left) time series of x, y, v (right) reconstruction results of v and q where each circle represents sampled time points (*n* = 4, M = 25, *N* =12, s = 9).

*where h*_*act*_, h_*ihb*_ *represents Hill functions for activation and inhibition respectively, and v_i_ represents arbitrary corruption shown in Figure 8 (a) and assumed to be sparse (at each time step, we choose card(v(j*)) = 1). *The magnitude of v_i_ are about 50% of the magnitude of 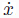. Since we consider arbitrary corruption, we need more time points (M > N). By using two-step refinements, first we estimate sparse large corruption as shown in Figure 8 (b) (top) and then, we reconstruct* **q**.

In practice, a specific node is corrupted by a hidden node and a small portion of the dataset can be largely corrupted by human error. Also, since we choose the set of possible candidate basis functions of the sensing matrix in (6), the columns of the sensing matrix may not be able to represent the underlying system (i.e., missing dictionaries). Then, we can consider the influence from these missing dictionaries as **v**.

**Figure 9.**
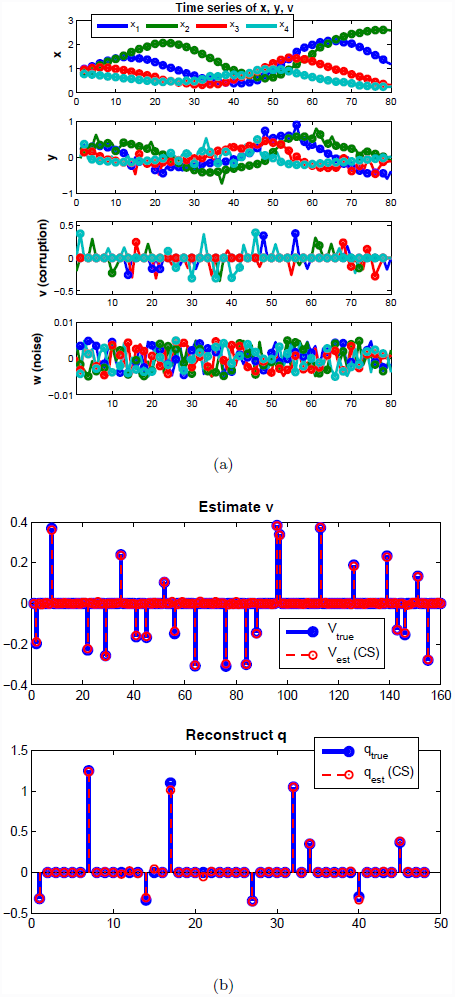
Arbitrary corruption with measurement noise. (Example 8) Reconstruction with corrupted signal. (left) time series of x, y, v (right) reconstruction results of v and q where each circle represents sample time points (*n* = 4, *M*= 40, *N* = 12, *s* = 9).

##### Example 8.

*(Arbitrary corruption with measurement noise) Recall a model (18) with different parameters and consider sparse large corruption* v *and small magnitude noise* w *(1% of the magnitude of 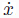). Figure 9 shows the time series data and reconstruction result*.

#### 3.3 Geometric view

In equation (17), since we assume all nodes are accessible and perfect measurement (meaning that there is no hidden node, v = 0 and no measurement noise, w = 0), we can solve equation (27) directly without filtering out the unmodelled dynamics (26) (i.e., 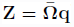). If there exist hidden nodes or measurement noise, we can still provide an unambiguous indication of the existence of these corruption (e≠ 0).

**Figure 10.**
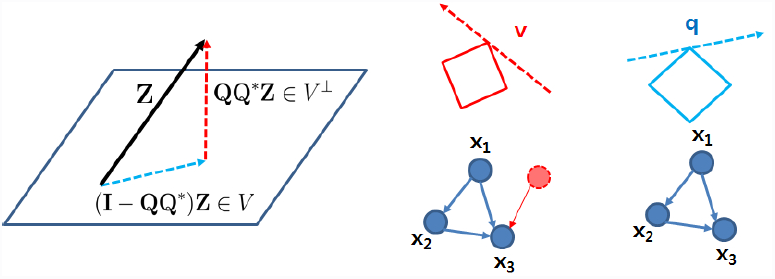
Geometric view of two-step refinement. A geometric view of two consecutive *l*_1_-norm optimizations.

The intuition is that 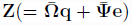 can be decomposed as the superposition of an arbitrary element in V(= Z − QQ*e) and of an element in V^⊥^(= QQ^*^ e) as shown in Figure 10. In other words, Z can be decomposed as the superposition of modelled dynamics and anomalies caused by hidden node or unmodelled dynamics. This geometric view enables us to understand how we could reveal deficiencies in our model:

- 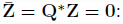 there is no hidden node ⇒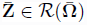
- 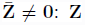 cannot be represented by 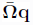 so there might be hidden nodes or our dictionaries in the sensing matrix 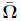 are not sufficient to represent Z (revealing deficiencies in our model or dictionaries).

### 4. HER2 Overexpressed Breast Cancer

We apply the proposed algorithm to study a breast cancer signaling pathway by reconstructing the graph structure using an RPPA dataset [27] as shown in Figure 11 (see also Figure S1, S2, and S3 for details: each figure presents the RPPA dataset and the result of graph reconstruction compared with *L*1, *L*2-optimization). Here, we choose small networks which are composed of 3 nodes and known to be sparsely connected, i.e., PI3K → PDK → Akt and PDK → Akt → mTOR in order to satisfy our assumption such that the influence on observable nodes from a hidden node should be sparse (i.e., **v** is sparse). The graph structures identified by the proposed method are consistent with the current understanding of the networks, whereas those found using *L*1- and *L*2-optimizations fail to reconstruct the known structure as shown in Figure 12.

**Figure 11.**
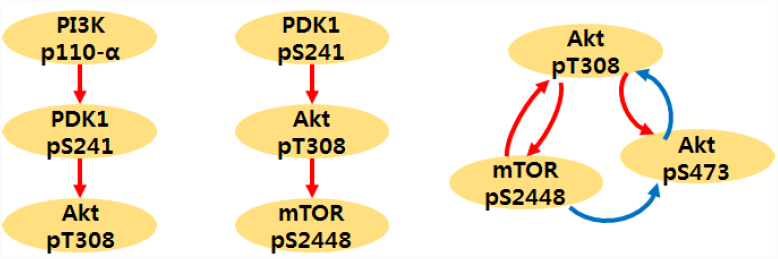
HER2+ overexpressed breast cancer. *CS* reconstruction result using Reverse Phase Protein Array data (`SKBR3` cell line, Serum [27]).See Figure S1, S2 and S3 for further details (red: activation, blue: inhibition).

**Figure 12.**
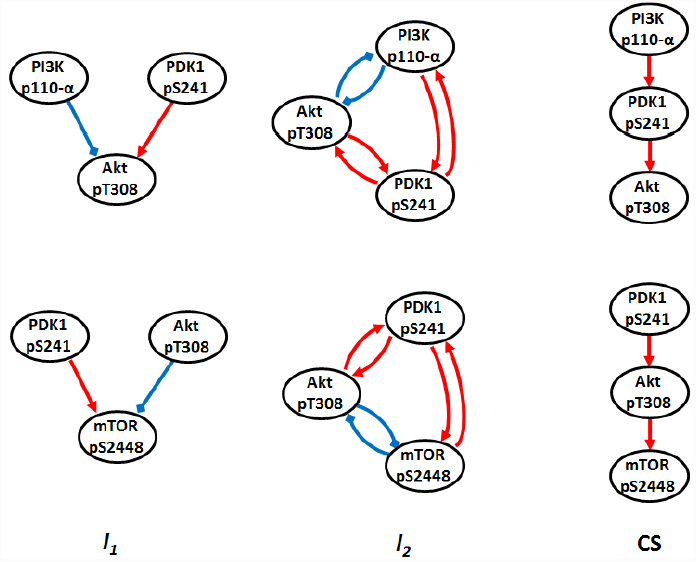
Sub networks inferred for the HER2/3 signaling network from Reverse Phase Protein Array data. The columns show the networks inferred by *L*1-optimization, *L*2-optimization, and *CS*. The network structures identified by *CS* agree with the current understanding of the network, whereas those found using *L*1 and *L*2 optimization do not.See Figure S1, S2 and S3 for further details (red: activation, blue: inhibition).

Also, an abstract model of the breast cancer signal pathway proposed by M. Moasser [31] is considered, as shown in Figure S3(b) where onospace?`PHLPP` isoforms are a pair of protein phosphatases, `PHLPP1` and `PHLPP2`, which are important regulators of Akt serine-threonine kinases (`Aktl`, `Akt2`, `Akt3`) and conventional protein kinase C (`PKC`) isoforms. `PHLPP` may act as a tumor suppressor in several types of cancer due to its ability to block growth factor-induced signaling in cancer cells [32]. `PHLPP` dephosphorylates Ser473 (the hydrophobic motif) in `Akt`, thus partially inactivating the kinase [33]. Unfortunately, in our `RPPA` dataset, we do not have PHLPP so we simply consider three nodes (`Akt`_pT308_, `Akt`_pS473_ and `mTOR`). Figure 11(right) shows the result of the proposed method using the RPPA dataset. The reconstructed graph structure matches up to known structure (Figure S3 (b)). Specifically, our result can capture the partial inactivating characteristics of PHLPP (i.e., mTOR(→ PHLPP) ⊣ Akt_pS473_).

## Conclusion

We proposed a method for reconstructing sparse graph structures based on time series gene expression data without any *a priori* information. We demonstrated that the proposed method can reconstruct graph structure reliably. Also, we illustrated that coherence in the sensing matrix can be used as a guideline for designing effective experiments.

Second, the proposed method is extended to the cases in which dynamics is corrupted by hidden nodes and the measurement is corrupted by human error in addition to the measurement noise. Using a two-step refinement procedure, we demonstrate good performance for the reconstruction of graph structure. A set of numerical examples is implemented to illustrate the method and its performance. Also, a biological example of HER2 overexpressed breast cancer using an RPPA dataset is studied. We are currently applying our method to recover the HER2 signaling pathway, where a significant part of the network is currently unknown.

## Method

### 1. Reducing Coherence by Transformation

#### 1.1. Rearranging the sensing matrix

Define z(*j*) as a vector of each component of y_*i*_ at the *j-*th time point:

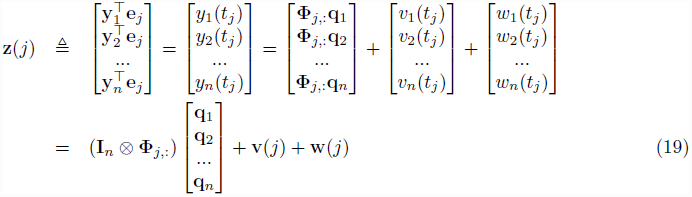

where Φ_*j*,:_ represents the *j*-th row of Φ. Consider 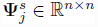 and multiply equation (19) by 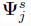:

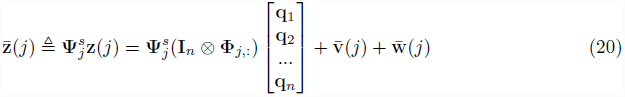

where 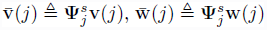 and rank 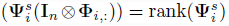 since rank 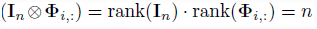. By stacking 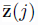,

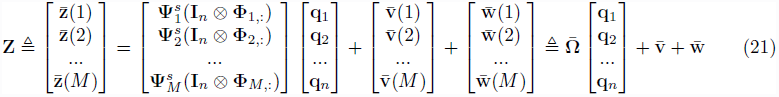

#### 1.2. Randomly chosen matrix Ψ

The optimization problem (14) is not trivial because of the constraint. One simple and heuristic approach is that we select 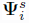 by (normalized) randomly chosen matrix with independent identically distributed (i.i.d.) random variable where 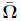 has full row rank, calculate 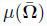 and run this with several times to reduce coherence. Since the randomly chosen matrix, 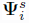, spreads out the component of Φ_*i*,:_ uniformly, we can reduce coherence.

#### 1.3. Finding the optimal transformation

This heuristic approach may not be enough to reduce the coherence. Consider a nonsingular matrix *P* ∈ ℝ ^*M*^ *^.n×M.n^* and 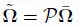 where 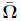 is constructed by heuristic way, i.e., randomly chosen matrix.

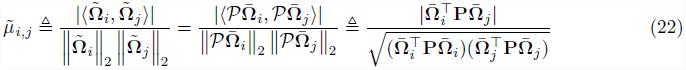

where 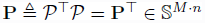 is positive definite and 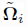 denote the *i*-th column of 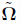. Note that if 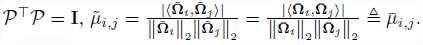. Therefore, our goal is finding P(= *P*^T^*P*) ∈ S^+^ such that

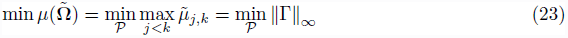

where Γ ∈ ℝ(*n.N*)^C2^ can be defined as follows:

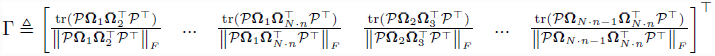

In practice, we ignore the denominator and solve the following problem:

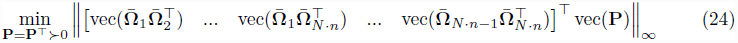

We can also combine ||•||_∞_ and ||•||_1_ to reduce the coherence. Note that for ||•||_∞_, we minimize the maxim coherence and for ||•||_1_, we minimize the sum of the all possible combinations of the columns of 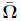, i.e., 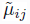. Thus, if certain bases are highly correlated, P or *P* makes the components of the sensing matrix spread out enough to differentiate the influences from those bases. Since we ignore the denominator, the optimal solution of (24) may be suboptimal. Thus, we can also combine heuristic way and (24) iteratively to reduce coherence of the sensing matrix in practice.

### 2. Two-step refinements

#### 2.1. Sparse large corruption with no measurement noise

Recall equation (13) where 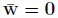:

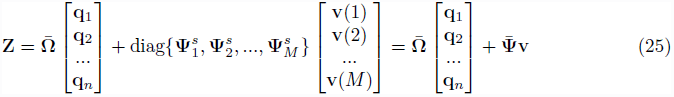

where 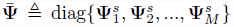 and v(*j*) represents sparse large corruption at the *j*-th time point, that could result from the existence of hidden nodes. We assume that the influence from hidden nodes is sparse and unknown (i.e., v(*j*) is assumed to be sparse). In other words, hidden nodes can affect only a few nodes’ dynamics (intuitively, if hidden nodes affect all nodes, there is no way to reconstruct the graph structure). Then, we consider the reconstruction of graph structure q from the corrupted signal Z.

Since we assume the number of rows of the sensing matrix 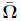 is greater than the number of columns *(M • n > N • n*), we consider Q^*^ which annihilates the sensing matrix 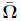 on the left 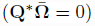 where Q^*^ is any (*M • n-N.n*)×*M.n* matrix whose kernel is the range of 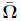 in ℝ^*M.n*^(*rank*(Q*) + *nullity*(Q^*^) = *M • n*):

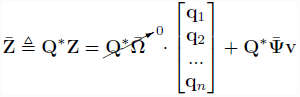

Then, the following two-step optimization problem enables us to compute q:

- Filter unmodelled dynamics out from the measurement:

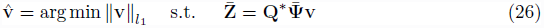
- Reconstruct q:

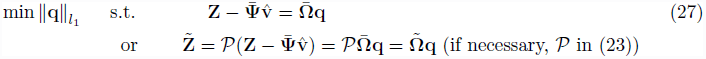

If we could somehow get an accurate estimate 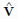 from equation (26), equation (27) represents the problem of reconstructing the graph structure q. The intuition is that 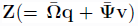 can be decomposed as the superposition of modelled dynamics and anomalies caused by hidden node or unmodelled dynamics.

The two step convex optimization problems (26) and (27) are *l*_1_-norm optimization problems in CS. Thus, if the sensing matrix 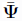 and 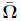 (or *P*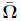) satisfy the incoherence condition, signals v and q can be recovered exactly [17] [29]. Here, there are many possible choices of Q* but we have to choose Q* to satisfy the incoherence condition for the exact recovery of v [17]. To choose such a Q^*^, we observe that 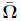 can be denoted as follows using Singular Value Decomposition (SVD):

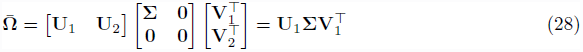

where 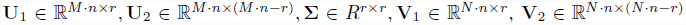 and *r* is the rank of 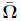.Suppose we choose Q^*^ such that 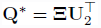.Then:

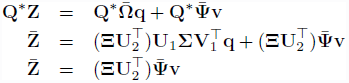

where **Ξ** can be used as a tuning matrix for satisfying the incoherence condition. A geometric view in **Results and Discussion: Section 3.3.** enables us to understand how we could reveal deficiencies in our model.

#### 2.2. Sparse large corruption with measurement noise

While considering influence from hidden nodes is interesting, it still may not be realistic to assume that except for hidden nodes, one is able to measure the node dynamics with infinite precision. A better model would assume that there is measurement noise. Consider the problem of recovering the graph structure **q** from the vector **Z** which is corrupted by measurement noise 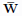 in equation (13):

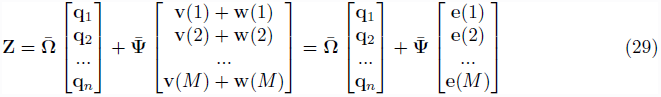

where **e**(*j*) = **v***(j*) + **w**(*j*), **w** is Gaussian noise *N*(0,σ) assumed to be bounded ||**w**||*_l2_* ≤ ϵ. In general, we can consider any corruption decomposed into sparse large error **v** and small magnitude error **w**[26]. Then, modified two-step refinements can be applied as follows:

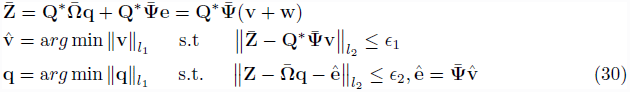

where the parameters ∈_1_,∈_2_ above depend on the magnitude of the small errors ∈, which can be determined as in [26].

## Author’s contributions

Conceived and developed computational method: YHC CJT, Analyzed the data: YHC CJT,Supervised the project: JWG CJT,Wrote the paper: YHC CJT

## Acknowledgements

This research was supported by the NIH NCI under the ICBP and PS-OC programs (5U54CA112970-08), and by the Stand Up To Cancer-American Association for Cancer Research Dream Team Translational Cancer Research Grant SU2C-AACR-DT0409 to JWG.

## Footnotes

1 If **A** is an *m* × *n* matrix and **B** is a *p* × *q* matrix, then the Kronecker product product **A** ⊗ **B** is the *mp*x*nq* block matrix: 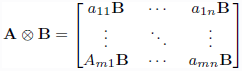

2 We compare the performance of CS with the performance of the L1 and L2 optimization as follows: 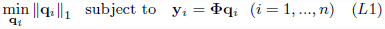 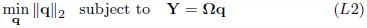

## Supporting Information

**Figure S1.**
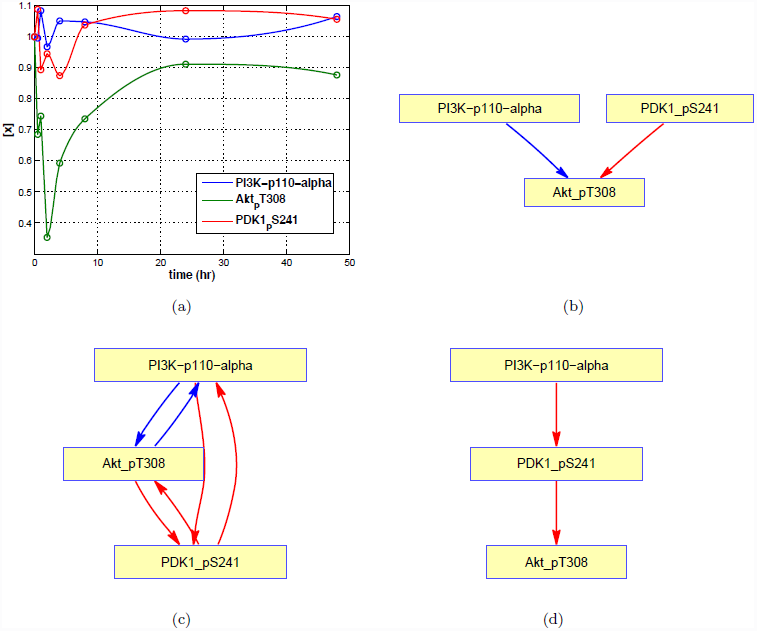
HER2+ overexpressed breast cancer. RPPA dataset (`SKBR3` cell line, Serum [27]) (a) gene expression data [0-48hr] (b) *L*1 optimization result (c) *L*2 optimization result (d) *CS* reconstruction result.

**Figure S2.**
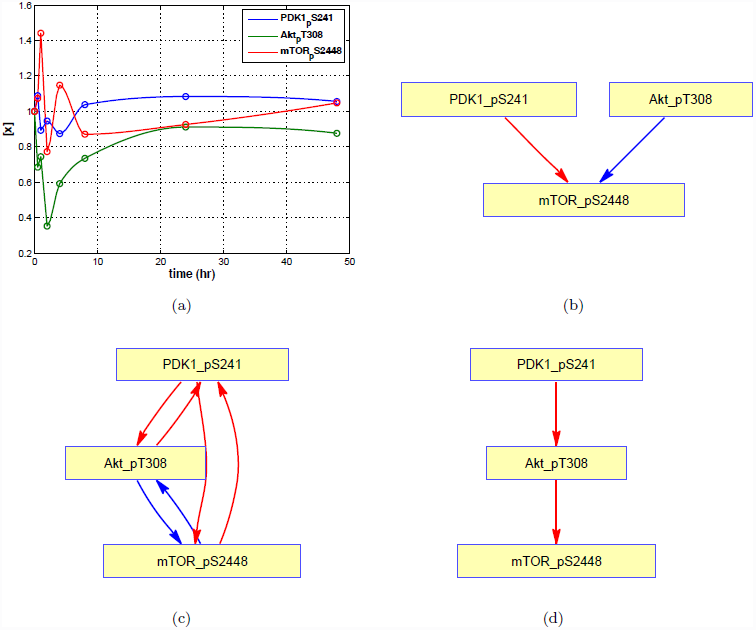
HER2+ overexpressed breast cancer. RPPA dataset (`SKBR3` cell line, Serum [27]) (a) gene expression data [0-48hr] (b) *L*1 optimization result (c) *L*2 optimization result (d) *CS* reconstruction result.

**Figure S3.**
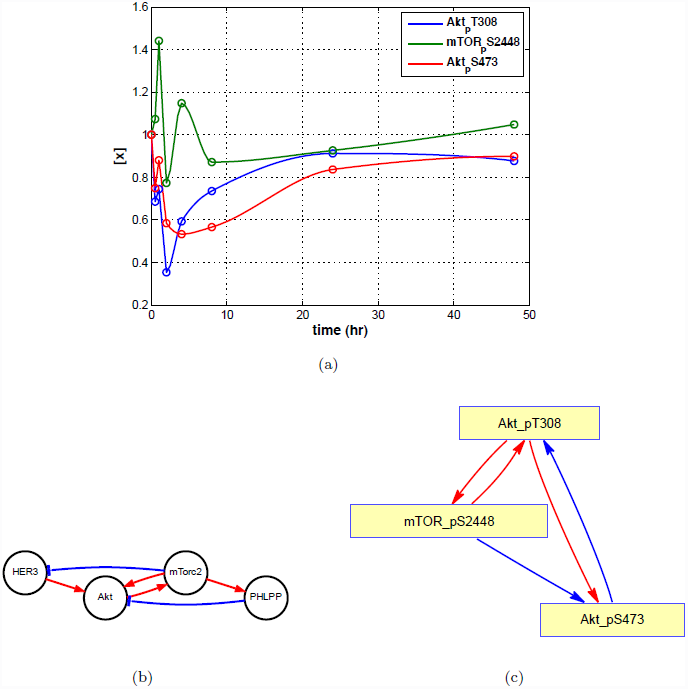
HER2+ overexpressed breast cancer. RPPA dataset (`SKBR3` cell line, Serum [27]) (a) gene expression data [0-48hr] (b) abstract model by Dr. Moasser (c) *CS* reconstruction result.

